# Relative humidity predominantly determines long-term biocrust-forming lichen survival under climate change

**DOI:** 10.1101/2020.06.09.141564

**Authors:** Selina Baldauf, Philipp Porada, José Raggio, Fernando T. Maestre, Britta Tietjen

## Abstract

1. Manipulative experiments show a decrease in dryland biological soil crust cover and altered species composition under climate change. However, the underlying mechanisms are not fully understood, and long-term interacting effects of different drivers are largely unknown due to the short-term nature of the studies conducted so far.
2. We addressed this gap and successfully parameterized a process-based model for the biocrust-forming lichen *Diploschistes diacapsis* as a common and globally distributed representative of biocrust communities to quantify how changing atmospheric CO_2_, temperature, rainfall amount and relative humidity affect its photosynthetic activity and cover. We also mimicked a long-term manipulative climate change experiment to understand the mechanisms underlying observed patterns in the field.
3. The model reproduced observed experimental findings: warming reduced lichen cover whereas less rainfall had no effect. This warming effect was caused by the associated decrease in relative humidity and non-rainfall water inputs as major water sources for lichens. Warming alone, however, increased cover because higher temperatures promoted photosynthesis during the cool morning hours with high lichen activity. When combined, climate variables showed non-additive effects on lichen cover, and fertilization effects of CO_2_ leveled off with decreasing levels of relative humidity.
4. *Synthesis.* Our results show that a decrease in relative humidity, rather than an increase in temperature may be the key factor for the survival of dryland lichens under climate change and that CO_2_ fertilization effects might be offset by a reduction in non-rainfall water inputs in the future. Because of a global trend towards warmer and thus drier air, this will affect lichen-dominated dryland biocrust communities and their role in regulating ecosystem functions, worldwide.

## Introduction

Biocrusts, communities dominated by lichens, cyanobacteria and mosses living on the soil surface, are a major biotic community in global drylands (Weber, Büdel, & Belnap, 2016). It is estimated that biocrusts cover around 12% of the land surface (Rodriguez-Caballero et al., 2018), providing important ecosystem functions across spatial scales. Locally, they prevent soil erosion (e.g. Bowker, Belnap, Bala Chaudhary, & Johnson, 2008; Cantón et al., 2011), enhance soil fertility by fixing atmospheric nitrogen (Barger et al., 2013; Xiao & Veste, 2017; Ferrenberg et al., 2018) and impact vascular plant species performance depending on their species composition (Havrilla et al., 2019) by for example inhibiting the germination of exotic plant species (Slate, Callaway, & Pearson, 2019). Globally, they contribute to the carbon cycle, directly by their own photosynthetic and respiratory activity (Porada, Weber, Elbert, Pöschl, & Kleidon, 2013; Rodriguez-Caballero et al., 2018) and indirectly by supporting carbon sequestration by vascular plants through nitrogen fixation (Weber et al., 2012). Due to their poikilohydric nature, biocrust constituents like lichens are highly adapted to high temperatures and limited water availability (Green, Sancho, & Pintado, 2011). However, empirical upscaling shows that climate and land use change will decrease their suitable habitat by up to 40% in the next decades, with semi-arid regions being among the most affected (Rodriguez-Caballero et al., 2018). If biocrusts are lost, the ecosystem functions they provide will change as well, for example leading to reduced hydrological control and alterations in C and N cycling (García-Palacios et al., 2018; Lafuente, Berdugo, Guevara, Gozalo, & Maestre, 2018; Reed et al., 2012).

Growing evidence shows that climate change will affect biocrust communities worldwide (Reed, Delgado-Baquerizo, & Ferrenberg, 2019), albeit the effects might be highly area- and species-specific. For example, studies from the Colorado Plateau found that warming and altered precipitation frequency led to a strong decline in moss cover and a shift towards cyanobacterial dominance (Ferrenberg, Reed, & Belnap, 2015; Reed et al., 2012; Zelikova, Housman, Grote, Neher, & Belnap, 2012). In an experiment from central Spain, simulated warming led to a substantial reduction in total biocrust cover and species richness (Ladrón de Guevara et al., 2018). In contrast to the studies from the Colorado Plateau, the cover of mosses increased with warming, but this did not suffice to compensate the drastic reduction in lichen cover observed (Escolar, Martínez, Bowker, & Maestre, 2012; Ladrón de Guevara et al., 2018).

Until now, studies on the response of biocrust communities to climate change are still rare compared to vascular plants (Reed et al., 2019). Considering the temporal and spatial scales at which climate change is operating, experiments and field studies have some major limitations. First, they are restricted to relatively small research areas and short time scales: all experiments conducted to date have been running for 15 years or less (Dacal et al., 2020; Ferrenberg et al., 2015). Second, manipulative climate change treatments potentially introduce unintended side-effects that can influence the results (Carlyle, Fraser, & Turkington, 2011) and different manipulation methods can hamper the comparison between studies (Bokhorst et al., 2013; Klein, Harte, & Zhao, 2005; Ladrón de Guevara et al., 2018). Also, with experiments alone it is difficult to understand climate change impacts on individual physiological processes, and thus gaining a mechanistic understanding of how these impacts translate to the observed changes in growth, cover and composition at the community level.

Mechanistic simulation models are valuable complementary tools to empirical studies because they allow to make projections in time and space, which are difficult to make with experiments, and analyze the underlying physiological processes leading to the observed impacts of climate change on organisms (Pacifici et al., 2015). For vascular vegetation, the gap between empirical and modeling research has been approached in various studies investigating the effect of increasing atmospheric CO_2_ and associated climatic changes using dynamic global vegetation models (DGVMs, e.g. Friend et al., 2014; Kolby Smith et al., 2016; Randerson et al., 2009). More locally, modeling studies have addressed topics such as the response of plant composition and traits to different rainfall regimes and aridity in dryland ecosystems (e.g. Henzler, Weise, Enright, Zander, & Tietjen, 2018; Lohmann, Guo, & Tietjen, 2018; Schwinning & Ehleringer, 2001). However, only few mechanistic modeling approaches focusing on biocrusts and non-vascular vegetation have been conducted so far (Kim & Or, 2017; Porada et al., 2013). The use of these models offers great promise to advance our capacity to gain a mechanistic understanding and predict future changes in biocrust communities due to climate change. However, we are not aware of the existence of any species-specific physiological model of major biocrust constituents (such as lichens) that has been used to assess climate change effects on these key organisms in drylands.

Here we used a mechanistic model developed to simulate a large number of artificial lichen and bryophyte strategies (LiBry, Porada, Tamm, Kleidon, Pöschl, & Weber, 2019; Porada et al., 2013) to assess the dynamics of the common biocrust-forming lichen *Diploschistes diacapsis* (Ach.) Lumbsch at two different sites in Spain under simulated climate change. This species has a wide geographical distribution and is among the most frequent biocrust-forming lichens worldwide (Bowker et al., 2016). Empirical studies show that this species is particularly affected by increased temperatures (Escolar et al., 2012; Ladrón de Guevara et al., 2018). We parameterized and validated the LiBry model for the first time for a single species and assessed its climate sensitivity towards changes in atmospheric CO_2_ concentration, rainfall, temperature, and relative air humidity. We used this model to: i) shed light on the long-term effects of changes in single climate variables compared to those of their combined changes on the physiological processes and resulting biocrust cover, and ii) determine the mechanisms leading to the observed decline in lichen cover under experimental warming in the field (Ladrón de Guevara et al., 2018). We also discuss the possibilities and current limitations of process-based models for assessing climate change impacts on biocrusts, as their use is being advocated for this aim specifically and for improving our understanding of biocrust ecology generally (Ferrenberg, Tucker, & Reed, 2017).

## Materials and Methods

### Species and site description

*Diploschistes diacapsis* is a terricolous, crustose, greyish-white lichen with a 1 to 3 mm thick thallus (Fig. S1) that is mostly found on calcareous substrate in exposed habitats (Lumbsch, 1988). *D. diacapsis* has a global geographic distribution (Galun & Garty, 2001; Ghiloufi & Chaieb, 2018; Pant & Upreti, 1993; Rosentreter & Belnap, 2001).

We simulated the physiological performance of D. diacapsis at two sites in Southeastern and central Spain, respectively: El Cautivo (37°0’N, 2°26’W, 200 m a.s.l.) and Aranjuez (40°2’N, 3°32’W, 590 m a.s.l.). Both sites are characterized by a semi-arid Mediterranean climate, with a higher mean annual temperature (MAT) and lower mean annual precipitation (MAP) in El Cautivo (MAT: 18.5 °C, MAP: 250 mm) than in Aranjuez (MAT: 15 °C, MAP: 350 mm). The cover of vascular vegetation is less than 40% at both sites, and mainly characterized by a mosaic of grasses, shrubs, and annual plants. The interplant spaces are covered with bare soil or well-developed biocrusts dominated by lichens such as *D. diacapsis* (see Maestre et al. (2013) for species checklist). Detailed site descriptions of El Cautivo and Aranjuez are provided in Büdel et al. (2014) and Maestre et al. (2013), respectively.

### Model description

For the simulation of *D. diacapsis* we used LiBry, a mechanistic model that simulates lichens, bryophytes, terrestrial cyanobacteria and algae (Porada et al., 2019, 2013). A full description of the model can be found in Porada et al. (2019, 2013), therefore we only briefly describe it here. LiBry was developed to quantify the global carbon uptake of non-vascular vegetation driven by climate and environmental conditions. These processes are implemented similarly as in DGVMs (e.g. Cramer et al., 2001; Pavlick et al., 2013), but were adjusted for lichen- and bryophyte-specific properties, such as poikilohydry or the dependence of CO_2_ diffusivity on the water content. LiBry is driven by hourly local climate input and environmental conditions determining the photosynthetic rate (based on Farquhar & Von Caemmerer, 1982) and thus gross primary productivity (GPP) and respiration (Q_10_ relationship). Both photosynthesis and respiration depend on the water saturation of the lichen. It increases through rainfall, snowmelt, dew and unsaturated air at relatively high relative humidity and decreases through evaporation, which depends on the surface energy balance of the thallus. Water saturation is also a proxy for lichen activity, which linearly increases between the minimum saturation necessary for metabolic activation (sat_min_) and the saturation at maximum activity (sat_max_). Net primary productivity (NPP) is based on the difference between GPP and respiration and is reduced by a species-specific constant turnover rate before being translated to the actual growth rate using the specific thallus area. The cover change in each time step consists of the growth rate multiplied by the existing cover and reduced by a constant mortality due to disturbances (e.g. perturbation by rabbits (Eldridge et al., 2010)).

In past applications, LiBry simulated the processes described above for different physiological strategies. Each strategy is defined by a unique combination of parameter values for 15 physiological traits and thus represents one (theoretical or actual) species. LiBry has been applied to assess the contribution of lichens and bryophytes to global cycles of biogeochemistry and hydrology (Porada, Van Stan, & Kleidon, 2018; Porada et al., 2013; Porada, Weber, Elbert, Pöschl, & Kleidon, 2014). However, it has never been applied to analyze species-specific responses at local scales as we are doing here.

### Model parameterization

Most physiological model parameters could be derived from a study on the differences in functional ecology of a sun and shade population of *D. diacapsis* close to El Cautivo (Pintado, Sancho, Green, Blanquer, & Lázaro, 2005). These include water storage capacity, water saturation at maximum activity, thallus height, optimum temperature for photosynthesis, Q_10_ value of respiration, and reference maintenance respiration at 10 °C. We used the mean value of these parameters obtained from sun and shade populations. Lichen albedo was calculated from a reflectance curve of a light-colored lichen dominated biocrusts with a high proportion of *D. diacapsis* (Chamizo, Cantón, Lázaro, Solé-Benet, & Domingo, 2012).

No direct measurements were available for some of the parameters necessary to calculate the photosynthetic activity of *D. diacapsis* (molar carboxylation and oxygenation rate of Rubisco (V_C,max_,V_O,max_), enzyme activation energy of Michaelis-Menten-Constants K_C_ and K_O_, and of V_C,max_ and J_max_ (electron transport capacity), thallus CO_2_ diffusivity). These parameters were calibrated using data on the relationship between net photosynthesis (NP) and light intensity at different temperatures and the dependence of NP on thallus water saturation (Pintado et al., 2005). For doing so, the relevant model functions to calculate NP were isolated from the model and calibration was done by visually assessing the differences between measured and modeled values for different parameter combinations within their global possibility range (Porada et al., 2013). The parameter combination that best fit the light curves at different temperatures and the water curve were taken as input parameters to the model (Fig. 1, light curves for all temperatures are shown in Fig. S2).

**Figure 1:**
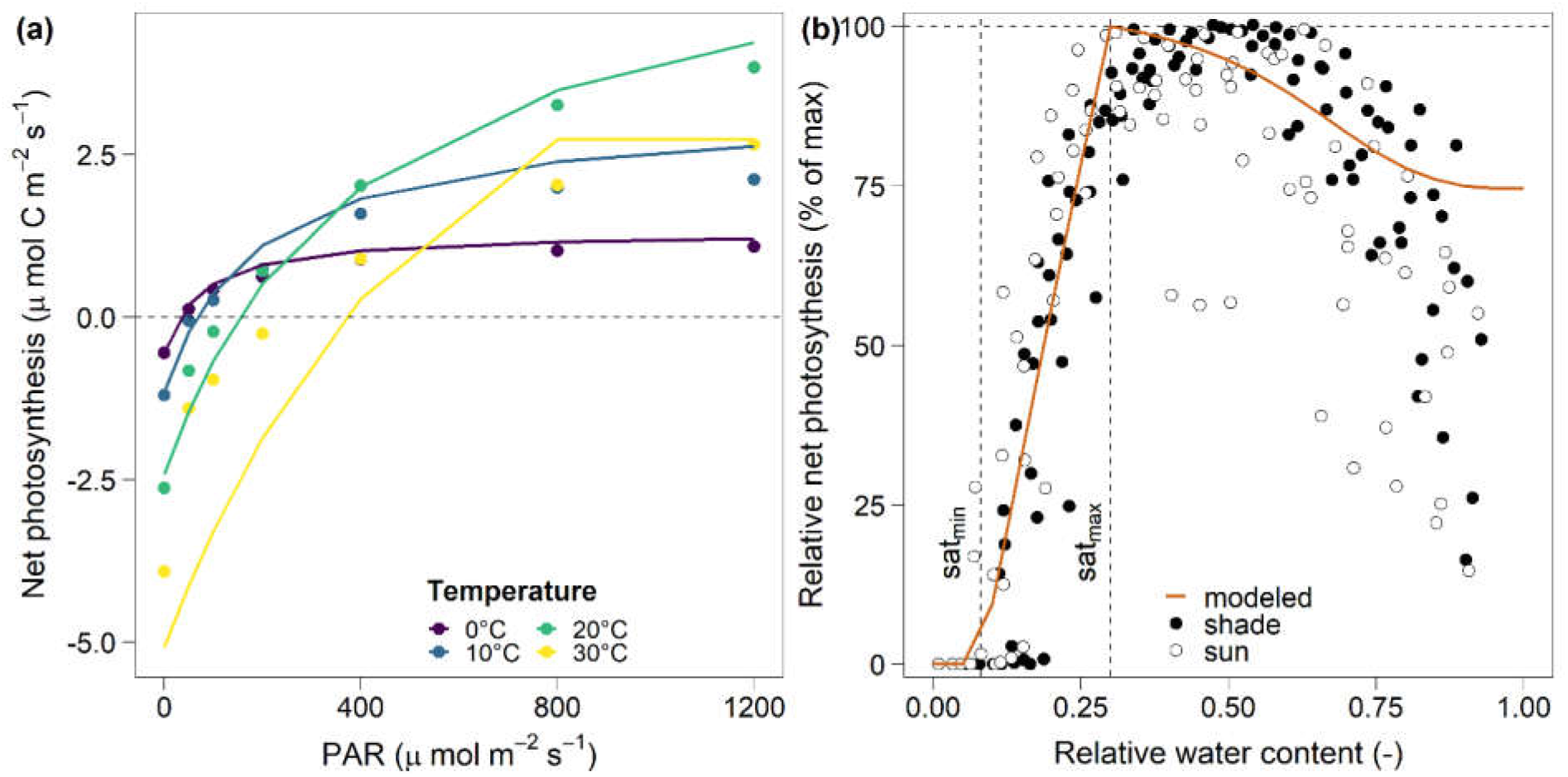
Calibration results for photosynthesis (data from Pintado et al., 2005). **(a):** light curves of net photosynthesis (NP) at different temperatures; points and lines represent field data (mean of sun and shade population) and modeled values, respectively. **(b):** water curve of relative NP at 15°C and 400 µmol m^-2^ s^-1^ light; points represent measured data for shade and sun populations. Dashed lines indicate the saturation needed for an onset of photosynthesis (sat_min_), and the saturation at which maximal photosynthesis is reached (sat_max_).

For the parameters thallus porosity and turnover rate, no values could be deduced from the available literature. Porosity was therefore assumed to be within a range of 0.3 and 0.4. Porosities lower than 0.3 led to specific thallus areas lower than 3.6 m^2^ thallus kg^-1^ C, which led to an unrealistic 100% mortality rate of *D. diacapsis* in El Cautivo. Porosities higher than 0.4 led to high specific thallus areas (>4.3 m^2^ thallus kg^-1^ C), which are unrealistic considering the structure of *D. diacapsis*. The constant turnover rate (how much of the lichen biomass is lost per time step) is unknown. Therefore, we assumed it to be 0.03 yr^−1^ which is at the lower end of the global value range reported by Porada et al. (2013) because of low turnover-rates expected in semi-arid environments.

In LiBry, the negative thallus water potential (Ψ_*H*2*O*_) increases with thallus water content and reaches zero at a specific water content *x*_ΦΘ,sat_, at which all water is stored extracellularly. Ψ_*H*2*O*_ influences how well the lichen can uptake water from the moisture in the air, which can contribute to metabolic activation. We could not find any data on Ψ_*H*2*O,min*_, *x*_ΦΘ,*sat*_, and the shape parameter *x*_Ψ*H2O*_ of the saturation dependent water potential curve for *D. diacapsis*. We therefore chose these values such that the obtained water potential curves are within the range observed for dryland lichens (Pintado & Sancho, 2002; Scheidegger, Schroeter, & Frey, 1995) (Fig. S3).

A more detailed description of the values of species parameters and the respective references is provided in Table S1. All simulations were conducted with a total of 22 strategies, each strategy representing *D. diacapsis* but with a different value for the unknown traits porosity (two possible values 0.3 and 0.4) and thallus saturation at which Ψ_*H*2*O*_ becomes negative (*x*_ΦΘ,*sat*_, eleven possible values from 0.05 to 1).

#### Modeling hydrophobicity of D. diacapsis

The thallus structure of *D. diacapsis* is characterized by a relatively impervious upper cortex, so it has a hydrophobic behavior when dry (Souza-Egipsy, Ascaso, & Sancho, 2002). This property can affect the lichen thallus itself (Pintado et al., 2005) and its impacts on infiltration (Cantón, Domingo, Solé-Benet, & Puigdefábregas, 2002). We therefore extended the original LiBry model and included a simple formulation of hydrophobicity by introducing the hydrophobicity factor f_*hyd*_, which reduces all water uptake by the lichen if the thallus is dry (i.e. if the thallus water content is below a critical saturation Θ_*crit*_ until which hydrophobicity occurs). The hydrophobicity factor f_*hyd*_ is calculated from two likely species-specific parameters: the critical saturation Θ_*crit*_, and *p*_*min,hyd*_ which is the minimum value for f_*hyd*_ at a water content Θ of zero:

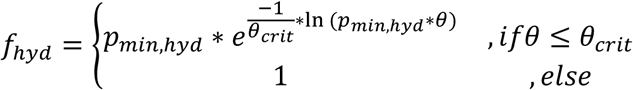

The hydrophobicity factor f_hyd_ increases exponentially, taking values from p_min,hyd_ for Θ = 0 to 1 for Θ = Θ_crit_ (Fig. S4). Modeled water uptake is then multiplied by f_hyd_. The parameters Θ_crit_ and p_min,hyd_ are unknown, and we could not find any indication in the literature as to how their values should be chosen. However, modeled lichen activity was sensitive to both parameters. We thus tested different value combinations and visually assessed the fit of modeled and measured lichen activity (Fig. S5). We selected the value combination of Θ_*crit =*_ 0.03 and *p*_*min,hyd*_ = 0.1 for all further analyses. Including this simple representation of hydrophobicity greatly improved the model fit of lichen activity to measured data from El Cautivo (Fig S5), indicating that hydrophobicity represents an important mechanism that should be considered when modeling biocrust lichens. Additional simulation results for the model without hydrophobicity and for a different value combination show quantitative but no qualitative differences, i.e. general model assessments are robust, despite the parameter uncertainty (Fig. S6).

### Climate forcing data

The model is driven by hourly climate data (long-wave radiation, short-wave radiation, snow cover, rainfall, relative air humidity, air temperature, and wind speed). For both simulation sites (El Cautivo and Aranjuez), we obtained on-site climate data of all necessary variables except longwave radiation, which was taken from the WATCH dataset (Weedon et al., 2011; see Porada et al., 2019 for detailed description of data preparation).

For El Cautivo, we used data from 2013 with a temporal resolution of 10 min, which were aggregated to hourly values (Büdel et al., 2014). Data gaps of more than four consecutive hours were filled with the values of the corresponding hours from the day before. Data gaps of less than four hours were linearly interpolated using the last and the next available data point. For the Aranjuez site, we used climate data from an *in situ* meteorological station (2009-2016). This dataset contained larger periods of missing data, which were filled with data from the nearby Aranjuez weather station of the Spanish meteorological service (AEMET), located 3,7 km away. If this data were not available, missing values were taken from 2012, a year without missing values. Due to the unreliable on-site wind measurements, wind data were taken from AEMET, and only larger gaps in the data were filled with field station measurements.

### Model validation

To validate the model results under current climatic conditions, we used chlorophyll fluorescence as an estimate of lichen metabolic activity and surface temperature field data of *D. diacapsis* from El Cautivo (Büdel et al., 2014; Raggio et al., 2017). These data were collected in 2013 and had a temporal resolution of 30 min. We compared daily and monthly activity and surface temperature patterns of simulated mean values in the steady state with measured chlorophyll fluorescence and temperature data.

### Simulation experiments

After model calibration and validation, we conducted two simulation experiments. The first one (hereafter Experiment 1) in El Cautivo, where we assessed the sensitivity of lichen physiological processes and cover towards systematic changes in single and combined climate variables. The second experiment (hereafter Experiment 2) mimicked and extended an ongoing climate change experiment in Aranjuez (Maestre et al., 2013) to determine the mechanisms leading to the observed changes in biocrust cover and to see how well the model can be transferred to other sites despite the intra-specific physiological variability of *D. diacapsis* (Lange, Belnap, Reichenberger, & Meyer, 1997; Pintado et al., 2005).

For Experiment 1, the initial lichen cover was set to 10%, and simulations were run for 900 years until a steady state in cover was reached. We first run the model under current climatic conditions with an atmospheric CO_2_ concentration of 395 ppm (values observed in 2013 (Tans & Keeling, 2020)). To test the sensitivity of *D. diacapsis* to altered climate conditions, we afterwards simulated two different climate change scenarios according to the Representative Concentration Pathway (RCP) 4.5 and 6.0 scenarios (Moss et al., 2010). For both scenarios, we altered climatic drivers (rainfall, temperature, air humidity) in isolation and combined, in each setting with the given concentration of atmospheric CO_2_ (RCP 4.5: 650 ppm, RCP 6.0: 850 ppm; Moss *et al.* 2010). The changes in climate variables were applied to hourly values, so that the annual variability in the time series remained the same. For the RCP 4.5 and 6.0 scenarios, we increased temperature by 3 °C and 5 °C, respectively, which corresponds to the projections of annual change in maximum temperature for Southern Spain in 2100 (Agencia Estatal de Meteorología (AEMET), 2020; IPCC, 2014). To keep the absolute amount of water in the air constant when increasing temperature, we scaled relative humidity down so that the actual saturation vapor pressure of the air remained the same across all temperature scenarios. Rainfall was decreased by 30% for both scenarios (Agencia Estatal de Meteorología (AEMET), 2020; IPCC, 2014).

Linear trend analyses showed that relative humidity in Southern Spain decreased between 0.7 and 2.5% per decade between 1973 and 2002 (Moratiel, Durán, & Snyder, 2010). We addressed this uncertainty by decreasing relative humidity by both 10% and 25% in each RCP scenario to represent two possible reductions in relative humidity over the next 100 years. In both RCP scenarios, we tested both interactions of all three variables. Additionally, we changed the same climate variables without increasing atmospheric CO_2_ in a control scenario.

For Experiment 2, the model was run with the exact same parameterization of *D. diacapsis* as for El Cautivo but driven by local climate resembling the ongoing climate manipulation experiment in Aranjuez (Escolar et al., 2012; Ladrón de Guevara et al., 2018). This full factorial experiment includes lichen-dominated biocrust plots under control and climate change treatments which are warming (on average +2.7 °C), rainfall exclusion (interception of 33% of rainfall) and a combination of both warming and rainfall exclusion (Fig. S7). Micro-climatic measurements showed that the warming treatments decreased relative air humidity (on average by 11.5% in 2016). For a detailed description of the experiment see Escolar et al. (2012).

We first generated new time series of temperature, humidity and rainfall mimicking the manipulation experiment in Aranjuez. For doing so, we used the biocrust surface temperature and near surface relative humidity measured within the experimental plots to calculate the relative difference of these two variables between control and warming treatments for each hour of the years 2016-2018. We applied the mean relative differences to the climate time series of Aranjuez to generate new time series of air temperature and relative humidity based on the observed differences between the treatments. For the months June, July, and August, no measurements were available, therefore we linearly interpolated hourly differences between May and September. To generate a time series with reduced rainfall, we reduced hourly rainfall values by 33%. With this new climate data, we conducted five simulation experiments: (i) control treatment without manipulations, (ii) rainfall exclusion, (iii) warming alone treatment with an increase in temperature (iv) warming treatment with an increase in temperature and a decrease in air humidity, and (v) a combination of (ii) and (iv). We ran the model for 900 years with an initial lichen cover equal to the mean initial cover value (68%) of all experimental plots. We then compared the changes in *D. diacapsis* cover driven by the treatments with the respective experimental results for biocrust-forming lichens. Since the modeled dynamics are slower than the observed ones, we compared the measured response over 10 years of climate manipulation with the steady state response in the model.

Since the model is deterministic, no replicates were simulated. In all scenarios of the two modeling experiments we simulated an undisturbed environment, in which *D. diacapsis* could potentially cover the full area.

## Results

### Model validation

In a steady state under current conditions, *D. diacapsis* was metabolically active 16% of the time during the year. In general, the model corresponded well with observed daily patterns of *D. diacapsis* activity in El Cautivo (Fig. 2, root mean square error (rmse) between 0.02 (June and July) and 0.26 (October)). Activity peaks occurred during the early morning hours, whereas activity during the day was very low. Most active hours occurred in autumn and winter (from September until January) and only a few where observed in the summer months (May until August). In September and October, the magnitude of the early morning activity peak was overestimated by the model, although the timing of the peak corresponded well to the measured data.

**Figure 2:**
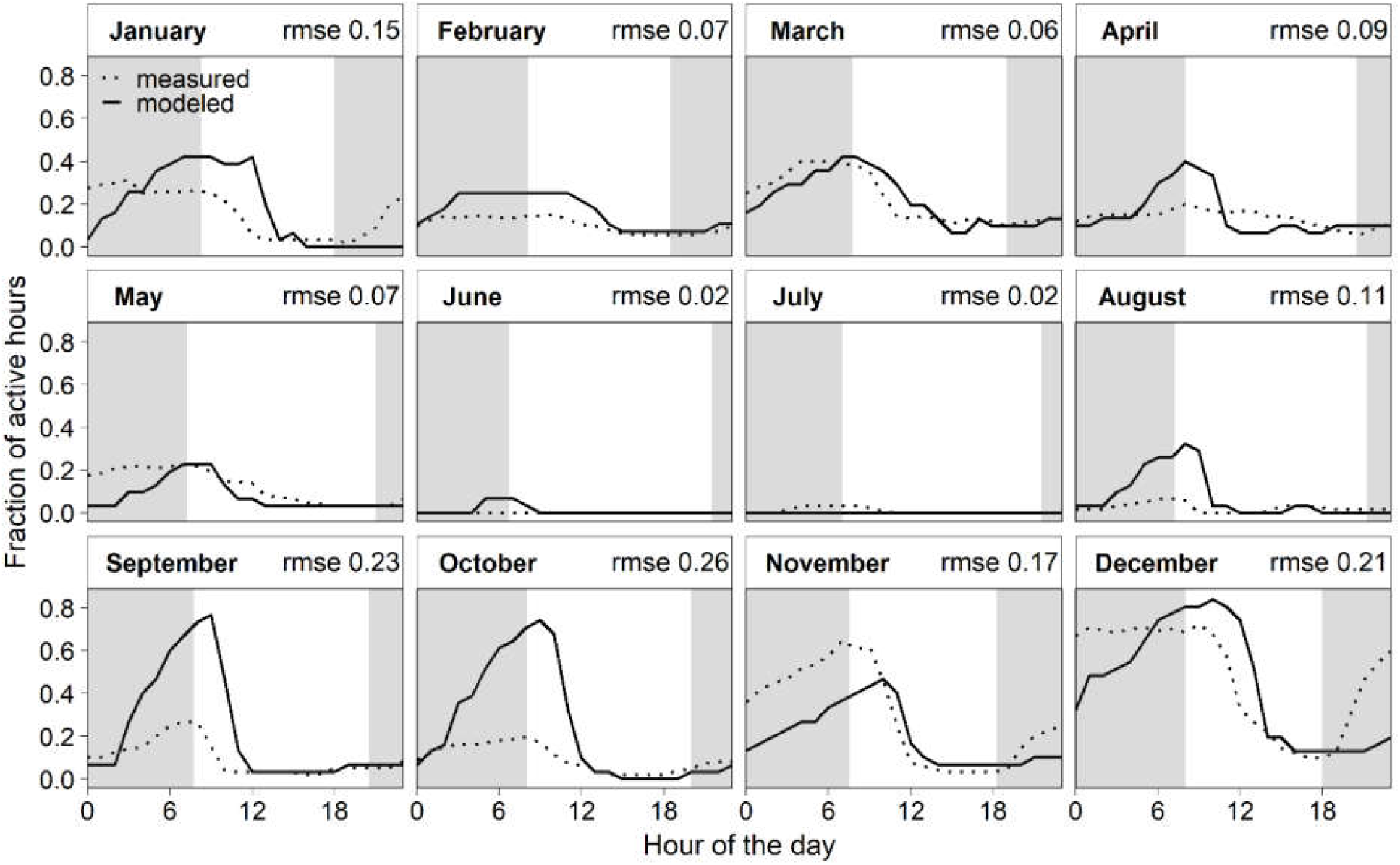
Daily fraction of active hours in each month. Modeled and measured values represent the mean of a binary representation of photosynthetic yield (yield/no yield; two samples) at the El Cautivo field site and binary values of activity represented by saturation status, respectively.

Average monthly thallus surface temperature was predicted very accurately, whereas maximum and minimum temperatures, respectively, were underestimated and slightly overestimated by the model (Fig. S8). The surface temperature peak in the warmer months was shifted by around two hours and underestimated in magnitude by the model (Fig. S9). However, during the most active hours of the day, the temperature validation was very satisfactory.

Simulated dew occurred during 78% of the nights and amounts to 30% of the time in a year. It was characterized by relatively small but constant watering events resulting in a total dew input of 24.8 mm yr^-1^, which was mainly accumulated during the nights between September and April and was particularly high in September and October (Fig. S10). Rainfall events occurred in 1.3% of the hours and generally delivered larger water amounts in a short period. Rain and dew impacted the thallus water saturation differently, as shown exemplarily for three days in Figure 3a,b. In most cases, rainfall led to an immediate increase in lichen water saturation to values exceeding the threshold saturation of 0.3 for maximum activity (sat_max_). In contrast, dew led to a more gradual increase in thallus saturation; a dew event must be sufficiently long and intense for the saturation to exceed the levels necessary for activation (sat_min_).

**Figure 3:**
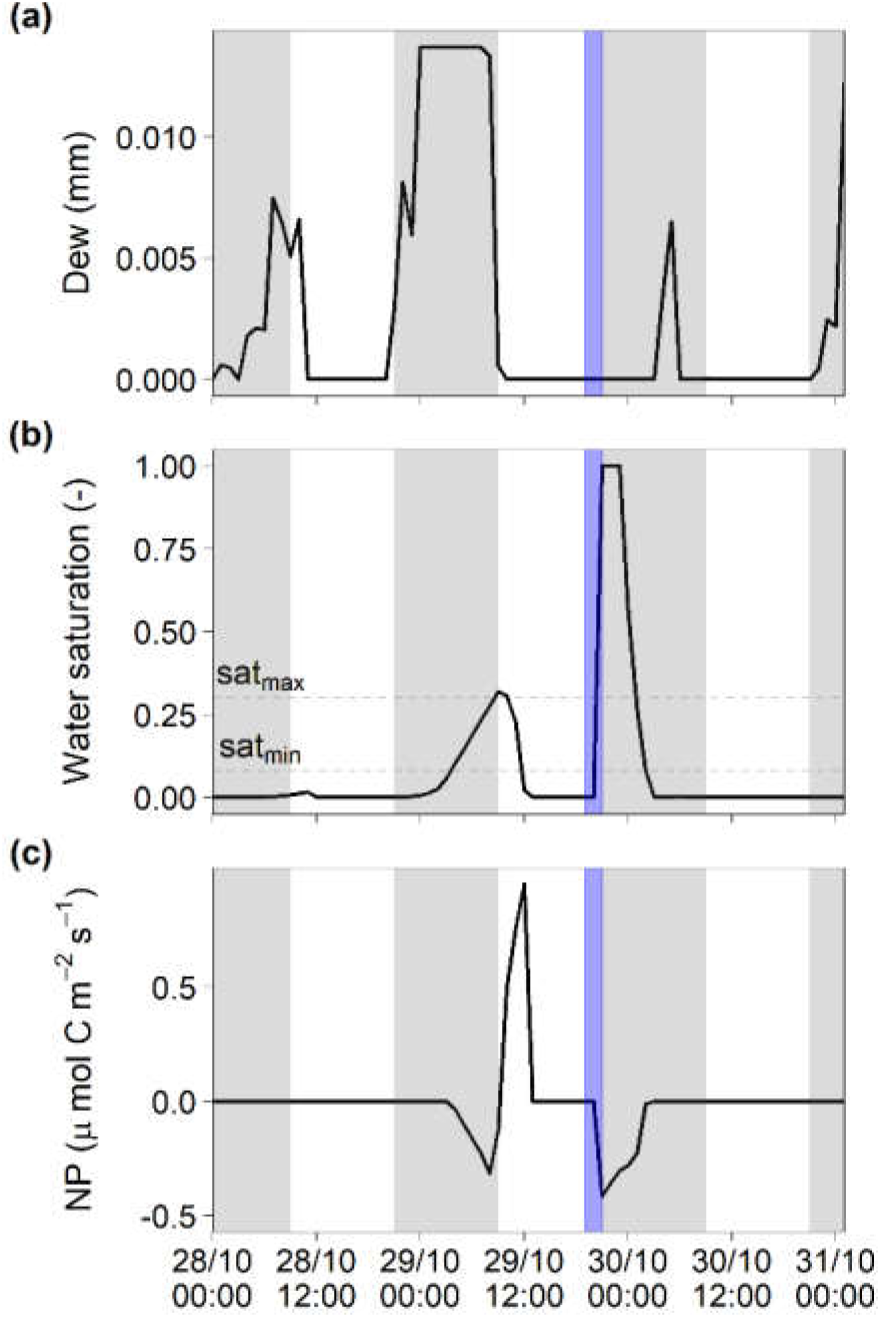
Modeled water saturation and net photosynthesis (NP) in response to dew formation and rainfall. **(a)**: simulated dew formation, **(b)**: water saturation with indication of saturation values of onset of photosynthesis (sat_min_) and maximal photosynthesis (sat_max_), and **(c)**: NP in hourly resolution for a period of two days in October 2013. Grey plot areas indicate nighttime, white areas daytime. The blue plot area marks a large precipitation event with a total amount of rainfall of 18.8 mm and a duration of 3 h.

### Experiment 1: sensitivity of lichen physiological processes and cover towards changes in climate

The modeled steady state cover of *D. diacapsis* under current climatic conditions was 37%. We found that, in the model, changing climatic variables had varying and interacting effects on the steady state cover (Fig. 4). We observed a consistent CO_2_ fertilization effect for all climate scenarios. Under current climatic conditions (control scenario) the net carbon gain was higher in the scenarios with increased CO_2_ despite the same annual activity time of 16% (annual NPP under current CO_2_: 8, RCP 4.5: 9.5, RCP 6.0: 10 g C m^-2^ s^-1^), leading to a cover increase by 30% and 32% for the RCP 4.5 and the RCP 6.0 scenario, respectively (Fig. 4). The cover difference between the control and RCP 4.5 scenarios was larger than the differences between RCP 4.5 and 6.0 scenarios, indicating a saturation type response of the CO_2_ fertilization effect. This was particularly visible under current climate and decreased rainfall.

**Figure 4:**
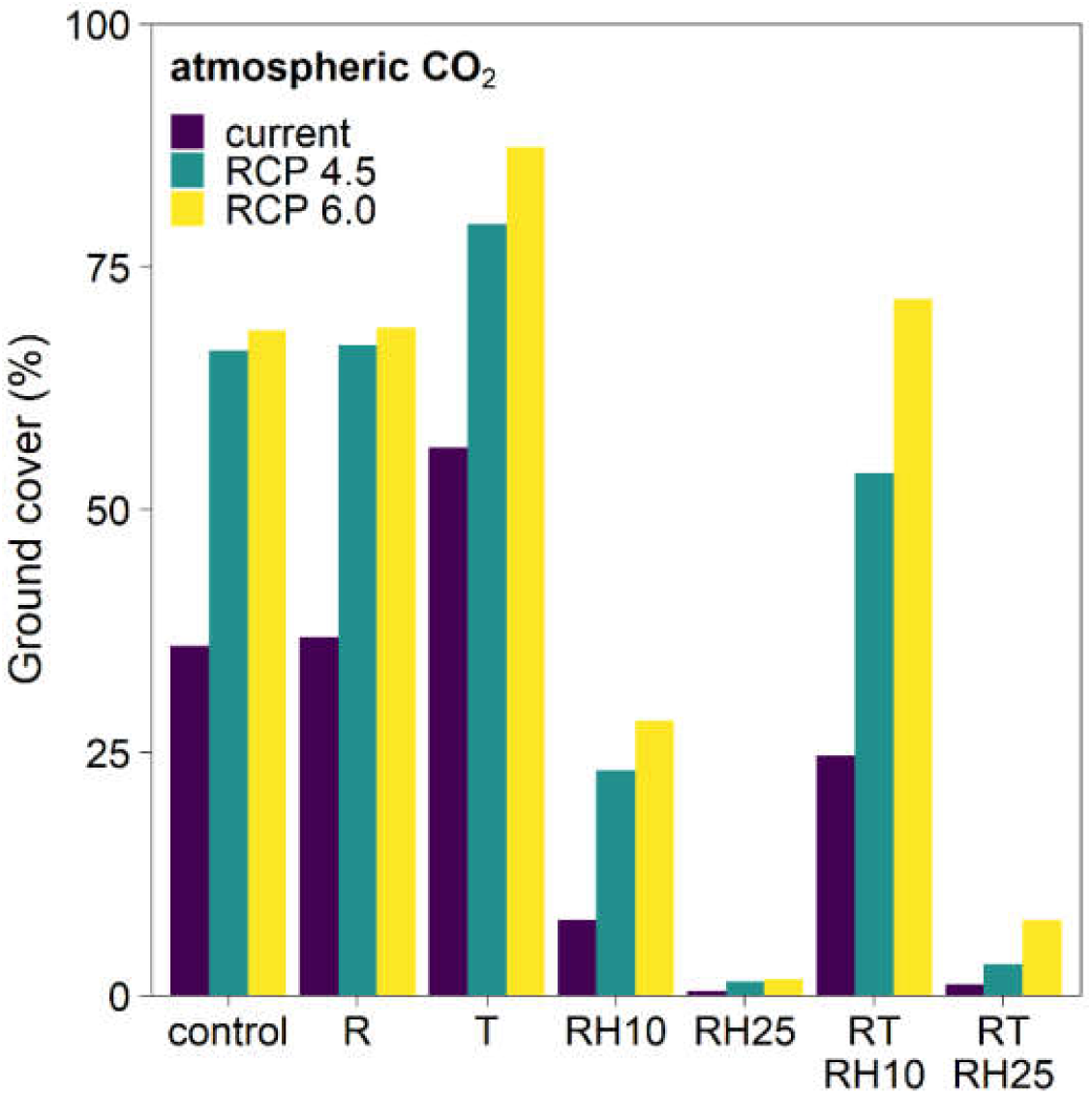
Steady state cover of *Diploschistes diacapsis* for different atmospheric CO_2_ (current = 395 ppm, RCP 4.5 = 650 ppm, RCP 6.0 = 850 ppm) and climate scenarios. Control = no changes in rainfall, temperature and relative humidity, R = rainfall reduction by 30%, T = temperature increase by 3 °C for current and RCP 4.5 scenario and 5 °C for RCP 6.0, RH10 = reduction of relative humidity by 10%, RH25 = reduction of relative humidity by 25%, RTRH10 = combination of R, T, and RH10 scenario, RTRH25 = combination of R, T, and RH25 scenario.

Increasing temperature by 3 °C or 5 °C had a positive effect on modeled steady state cover (Fig. 4). The activity between the three temperature scenarios was similar (annual active time fraction = 18%, 18% and 19% for current, RCP 4.5 and RCP 6.0 scenarios) because moisture availability was the same in all of them. A reduction in rainfall did not show an effect on lichen cover in any climate change scenario and climate variable combination.

A decrease in relative humidity by 10% and 25% led to a substantial decrease in lichen cover to values below those of the control scenario in both RCP scenarios considered (Fig. 4). At a 25% reduction of relative humidity, lichen cover was reduced to values from 1 to 8% depending on atmospheric CO_2_ resulting from a reduction in dew input (decrease by 31 to 36% relative to control) and associated activity time (decrease by 63 to 69% relative to control) (Table S2).

The climate sensitivity results indicate an interaction between the effects of the single climate variables because the cover change of the combined scenarios differed from the additive changes of the respective single variables. Generally, cover decline was lower for the combined scenario with 10% lower relative humidity (17% for RCP 4.5 and 24% for RCP 6.0) and higher for the one with a 25% lower relative humidity (11% for RCP 4.5 and 13% for RCP 6.0) compared to what would have been expected from adding the single effects.

### Experiment 2: Revealing the mechanisms leading to observed cover decline under climate change

Steady state cover of the control scenario in Aranjuez was higher than in El Cautivo (Fig. 5c). However, the observed effects of warming and rainfall reduction on the cover of *D. diacapsis* were qualitatively comparable between these sites (Fig. 5c). In the steady state, we found no effect of rainfall exclusion on lichen cover (55% cover for both control and rainfall exclusion treatments). Warming alone had a slight positive effect on steady state cover (59%) but the associated changes in relative humidity as observed in the field experiment led to a decrease in lichen steady state cover (warming scenario: 43%, warming and rainfall exclusion: 44%). The qualitative effects of climate change treatments on modeled cover were similar to the observed effects (Fig. 5a,b), but modeled effects were not as strong as those observed in the field.

**Figure 5:**
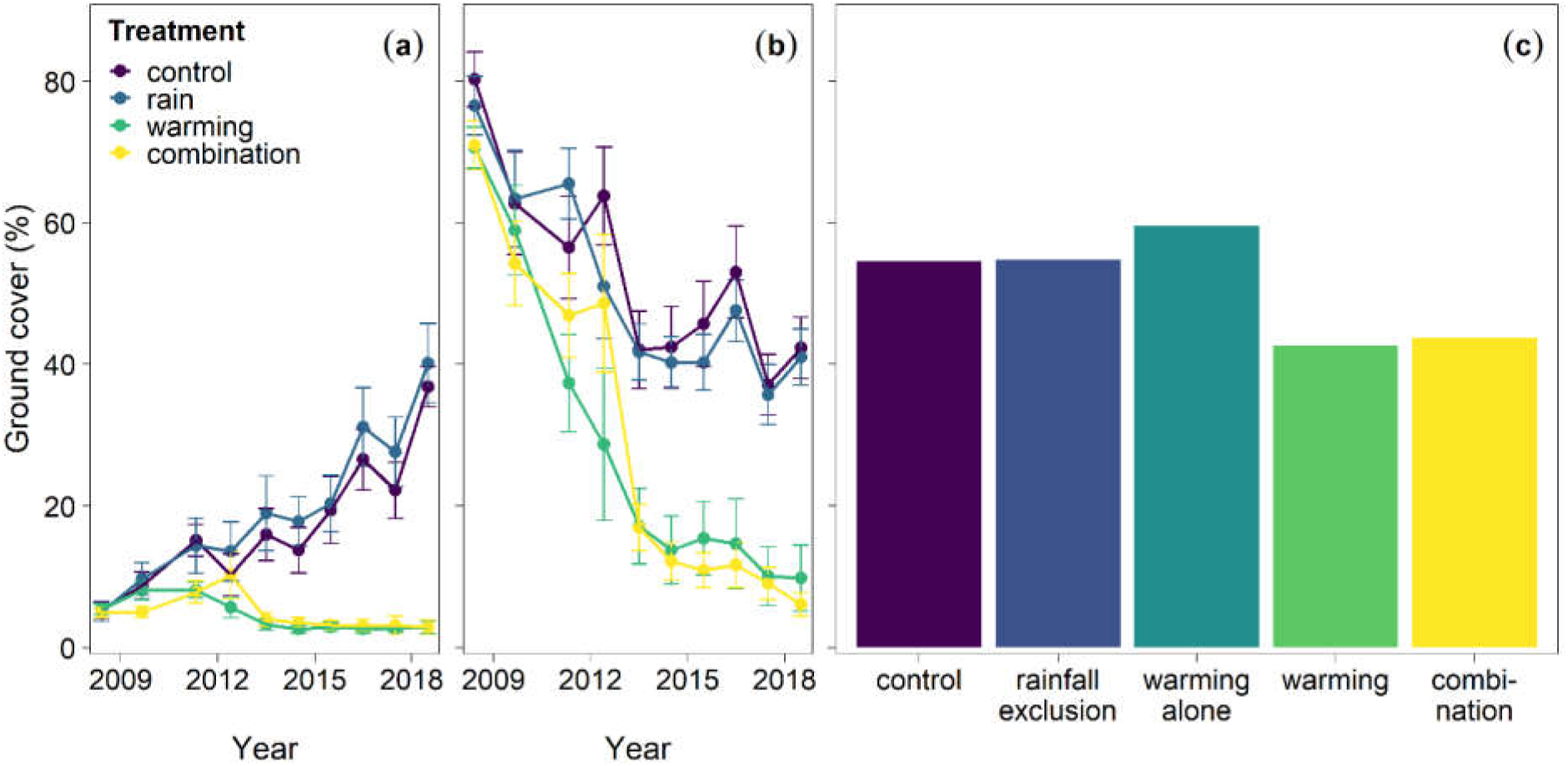
Changes in the cover of *Diploschistes diacapsis* as a result of climatic conditions in the Aranjuez experiment and in the model. **(a)**: measured cover change of biocrust lichens (including D. diacapsis) in plots with an initial cover < 10%, **(b)**: same as (a) but with an initial cover > 75%, and **(c):** modeled cover in steady state. Rainfall exclusion refers to a 30% reduction in rainfall, warming alone to an increased temperature by 2.7 °C on average, warming an increase in temperature by 2.7 °C and an associated reduction in relative humidity, combination is a combination of the scenarios rainfall exclusion and warming (associated with reduced relative humidity).

## Discussion

### LiBry reproduces physiological behavior and cover of *D. diacapsis*

Overall, the validation results were satisfactory; LiBry accurately predicted mean daily activity for all months except September and October and daily lichen surface temperature for the midday peak temperatures from April to October. During these hours, however, the lichen was mostly inactive and this difference between measured and modeled temperature should not influence modeled lichen.

Periods of activity over- and underestimation by the model can be explained by two different mechanisms. First, the underestimation of activity during some periods (especially November until January) could be explained by different mechanisms of rainfall activation in model and field. In the field, rainfall is an important source of hydration, leading to extensive moist periods in the winter months (Raggio et al., 2014). Rainfall water can be stored within the thallus, on its surface, or as soil moisture (Berdugo, Soliveres, & Maestre, 2014). This temporal storage can indirectly lead to longer activation periods through rainfall and are not represented in the model, where excess water is lost from the system. Second, the overestimation of activity could be explained by the thallus saturation model. The high activity in September and October can thus be explained by the relatively high dew input in these months (Fig. S10). One potential reason for this is that dew is estimated correctly by the model, but the dew amount taken up by the thallus is higher in the model compared to the field (either due to evaporation or underestimated hydrophobicity). Alternatively, the model may have overestimated dew inputs, and thus lichen activity. However, dew production in the model agrees with measured values from a nearby area (Cabo de Gata-Níjar national park), where dew occurs in 78% of the nights (Uclés, Villagarcía, Moro, Cantón, & Domingo, 2014). The estimated total amounts differ between that study (35-57 mm y^-1^ from 2007 to 2010) and our model (25 mm yr^-1^). Apart from the inter-annual and spatial variability in dewfall, these discrepancies might result from different reference surface areas. Model results reflect dew formation on the thallus surface, whereas the field study reflects a range of surface covers, including plants and stones that have higher relative contributions to dew formation compared to biocrusts (Uclés, Villagarcía, Cantón, & Domingo, 2016).

The analysis of physiological processes in hourly resolution (Fig. 3) showed that the diurnal response of modeled lichen hydration and NP is similar to the responses observed in the field. Dew activation during the night was the main hydration source in the model, leading to 89% of the lichen’s active time, whereas rainfall alone only accounted for 6% of active time. In 5% of the time there was an overlap when both watering events occurring simultaneously. The activity window caused by dew simulated by the model is typically longer than that caused by rain. This finding is not fully supported by field observations that showed the longest activation periods from rain events followed by cloudy days in the same research area (Green, Pintado, Raggio, & Sancho, 2018; Raggio et al., 2014). However, this does not contradict the well-known reliance of biocrust-forming lichens on dew in the study area, which allows for frequent net carbon gains independent of rainfall and interrupts long desiccation periods that negatively impact their physiological performance (del Prado & Sancho, 2007; Green et al., 2011; Pintado, Sancho, Blanquer, Green, & Lázaro, 2010; Raggio et al., 2014).

Modeled steady state cover under current conditions (37%) corresponds to the total biocrust cover in the Tabernas region (40-45%), but it is higher than the measured proportion of chlorolichens (15%) (Büdel et al., 2014). This discrepancy is not surprising, since the model did not include competition for space with vascular plants and other lichens and bryophytes, which is intense in these communities (Maestre, Escolar, Martínez, & Escudero, 2008). Against this background, the modeled steady state cover seems to be within a reasonable range for this ecosystem, and falls within what has been observed in the field (Lázaro et al., 2008).

### Relative humidity drives climate change responses of *D. diacapsis*

Although changes in single climate variables are unrealistic under climate change conditions, doing this using a simulation model can help to disentangle the overall effects of climate change and determine how the single variables interact, something not always possible to do with field experiments. Overall, the modeled response of *D. diacapsis* to changes in single climate variables was in good agreement with laboratory and field measurements.

Carbon exchange studies with biocrust-forming lichens (including *D. diacapsis*) show increasing photosynthetic rates with CO_2_ partial pressure, albeit the effect size varies between species and depends on the thallus water content (Lange, 2002; Lange et al., 1997; Lange, Green, & Reichenberger, 1999). However, it must be considered that modeled carbon uptake and growth are not nutrient limited because nitrogen and phosphorus cycles are not included in the model. In reality, the positive CO_2_ fertilization effect will probably be counteracted by limited nutrient availability (Goll et al., 2012). If the crust is active (i.e. no water limitation), modeled photosynthesis is generally light- and temperature-limited, which partly explains the small cover difference observed between the RCP 4.5 and 6.0 scenarios.

Increased temperatures resulted in a higher NP of *D. diacapsis* in the model. This lichen is mainly active in the early morning hours, when conditions for photosynthesis are suboptimal (low radiation and temperature, Fig. S11) and resemble more temperate environments (Pintado et al., 2010). Under these conditions, NP can benefit from higher temperatures given enough moisture from overnight dew.

Although both rainfall and non-rainfall water inputs (NRWI) are important sources of hydration for biocrust-forming lichens (Raggio et al., 2014; Raggio et al., 2017), our modeling results show no effect of a decrease in overall precipitation but a very large effect of decreases in relative humidity. These results underline the importance of NRWI for *D. diacapsis* (Pintado et al., 2010) because a reduction in relative humidity drastically reduces dew input and thus leads to a reduction in activity time and lichen cover. This corresponds to field results from Aranjuez showing a much larger influence of reductions of NWRI driven by experimental warming than of rainfall on the photosynthetic performance of biocrust-forming lichens (Ladrón de Guevara et al., 2014). The lack of effects of rainfall reduction on lichen cover can be explained by the fact that, albeit reduced, rainfall events are still large enough to saturate the lichen. Accordingly, other studies have shown that the timing, size and frequency of individual rainfall pulses, rather than average rainfall amount, affect biocrust performance and cover (Baldauf, Ladrón de Guevara, Maestre, & Tietjen, 2018; Belnap, Phillips, & Miller, 2004; Zelikova et al., 2012).

We found an interaction rather than an additive effect when changing all climate variables at the same time. Generally, our results suggest that an increase in atmospheric CO_2_ could mitigate some of the negative effects of reduced water availability, and that this effect is larger at higher temperatures. However, the net benefits of mitigation at higher atmospheric CO_2_ become smaller if relative humidity decreases (a trend being already observed in Spain, Moratiel et al., (2010)). Additionally, there is a general trend towards warmer and dryer soils that can further reduce water availability and increase drought stress for biocrusts (Soong, Phillips, Ledna, Koven, & Torn, 2020). Field studies on this subject are still very rare, but a study conducted in the Mojave desert suggests that higher atmospheric CO_2_ cannot mitigate the negative effects of drought on biocrust cover (Wertin, Phillips, Reed, & Belnap, 2012). Our results highlight the key role of relative humidity and although its importance for biocrust activity has been discussed in empirical studies (Pintado et al., 2010; Raggio et al., 2017), modeling allowed us for the first time to quantitatively and qualitatively compare its effect to those of other climate drivers. *D. diacapsis* is a biocrust-forming lichen with global distribution (Bowker et al., 2016) and trend analysis of climate data from 1973-2005 shows a decrease in relative humidity in many dryland regions worldwide (Willett et al., 2014). Therefore, the observed and simulated trend in biocrust cover decline at our study sites is likely to also represent a global decline in lichen-dominated biocrusts. For global vegetation, this association has already been shown with satellite-based models which suggest that a positive CO_2_ fertilization effect is offset by the increase in vapor pressure deficit (i.e. reduction in relative humidity) leading to an overall decrease in the NDVI (normalized difference vegetation index), leaf area index and estimated gross primary productivity (Yuan et al., 2019).

### Modeling results mimic observed responses in the field

Application of the model to the climate change experiment in Aranjuez showed similar effects as the systematic climate sensitivity analysis in El Cautivo. We found no effect of rainfall exclusion, and a negative effect of both warming and a combination of warming and rainfall exclusion on lichen cover. Warming alone, without the associated reduction in relative humidity, had a slightly positive effect.

Model results are in line with field observations, which showed no significant effect of decreased rainfall, but a strong negative response of lichen to increased temperatures. Between 2008 and 2011, *D. diacapsis* cover declined by around 8% in the warmed plots and by roughly 5% in the plots with both, warming and rainfall exclusion (Escolar et al., 2012). Total biocrust lichen and *D. diacapsis* cover continued to decline until the total cover difference between warmed and non-warmed plots was about 40% in 2016 (Ladrón de Guevara et al., 2018). The modeled steady state cover differences between warmed and non-warmed treatments (difference of ca. 10% cover) correspond well to the reported values for the first period of the experiment (Escolar et al., 2012). However, the model did not reproduce the drastic further decline in lichen cover over the next years. This is not surprising as Ladrón de Guevara et al. (2018) stated that the rapid loss of lichen cover could partly be explained by the easy detachment of the lichen thalli from the soil surface and the consecutive loss of thallus parts through wind. This process is not represented in the model; thus, it was to be expected that modeled cover losses are less drastic than the observed losses. Additionally, the physiological trait values of *D. diacapsis* are variable between different locations and even between plots of different exposure within one site (Lange et al., 1997; Pintado et al., 2005). Therefore, the population in Aranjuez might differ in some physiological parameters from the population in El Cautivo, which could further explain differences between model and experiment.

Escolar et al. (2012) hypothesized that the decline in cover under warming could be promoted by an associated increase in respiratory carbon losses, which could not be compensated by photosynthetic activity. However, they also found a consistently higher photosynthetic efficiency (measured as F_v_/F_m_ ratio) under warming and therefore raise doubts about this hypothesis. With the model, we showed that increasing temperatures alone led to an increase rather than a decline in lichen cover, which is consistent with their observation of a higher photosynthetic efficiency under warming. The indirect effects of warming on relative humidity and therefore NRWI were responsible for the cover decline in the model, therefore, we hypothesize that they were also responsible for the cover decline observed in the field experiment (Ladrón de Guevara et al., 2018).

## Conclusions

Our modeling results provide the first forecasts of long-term climate change effects on a dominant biocrust-forming lichen. They highlight the importance of relative humidity as a driver of the physiological responses of *D. diacapsis* to climate change and indicate that increasing CO_2_ concentrations could mitigate the effects of decreasing water availability to a certain degree. Negative effects of drier air rather than higher temperatures might be the key factor in determining dryland lichen survival and cover under future conditions. Global climate trends suggest that this mechanism is of relevance for many lichen-dominated dryland biocrust communities that rely on dew or relative humidity as a major water source. Our results highlight the value of process-based modeling to disentangle the effects and interactions of major climate change drivers acting simultaneously and in isolation, something not often possible to do in the field, and provide guidelines for future climate change experiments with biocrusts. They should explicitly consider the indirect effects of increased temperature on relative humidity and NRWI, especially in areas where those are important sources of biocrust hydration. We showed that a detailed understanding of the underlying processes by complementing experimental work with modeling is necessary to explain non-additive effects of altered climate drivers on biocrust performance. This will hold even more in more complex studies focusing on whole biocrust communities.

## Acknowledgements

This research was supported by the Collaborative Research Centre 973 (www.sfb973.de) of the German Research Foundation (DFG) and by the European Research Council (grant agreement nº 647038 (BIODESERT)). P. Porada appreciates funding by Deutsche Forschungsgemeinschaft (DFG, German Research Foundation) – 408092731. F. T. Maestre acknowledges support from Generalitat Valenciana (CIDEGENT/2018/041) and the Alexander Von Humboldt Foundation. J. Raggio acknowledges the ERA-Net BiodivERsA program as Soil Crust InterNational (SCIN) and The Spanish Ministerio de Economía y Competitividad (MINECO) project numbers PRI-PIMBDV-2011-0874 and CRYPTOCOVER, CTM2015-64728-C21-R. We thank Cristina Escolar, Victoria Ochoa, Beatriz Gozalo, Sergio Asensio, Roberto Lázaro, David Sánchez-Pescador, Mónica Ladrón de Guevara and Patricia Alonso for their help with the setting up, maintenance and monitoring of the field experiments. We thank Bettina Weber and Burkhard Büdel for providing the meteorological data for El Cautivo which was collected within the Soil Crust InterNational (SCIN) project.

## Author contributions

SB, PP, BT and FTM. planned and designed the research, SB conducted the modeling experiments, JR and FTM provided field data, SB and PP did the data analysis, all authors contributed to data interpretation, SB wrote the manuscript and all authors revised it.

## Notes

https://doi.org/10.6084/m9.figshare.c.4970051.v1

### Competing Interest Statement

The authors have declared no competing interest.

