## Supporting Information for "Relative humidity predominantly determines long-term biocrust-forming lichen survival under climate change"

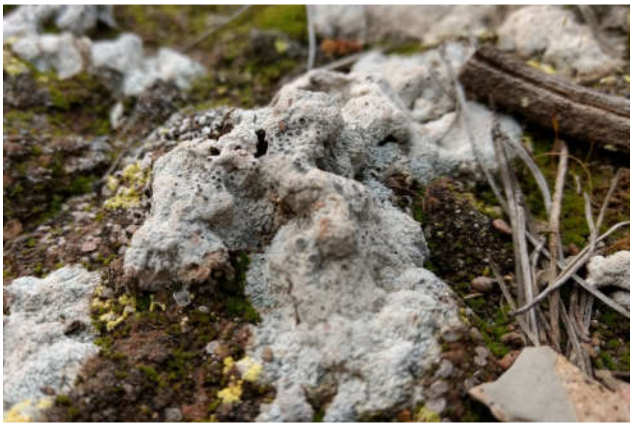

**Fig. S1** *Diploschistes diacapsis* dominated biocrust from Aranjuez (Photo by Fernando T. Maestre).

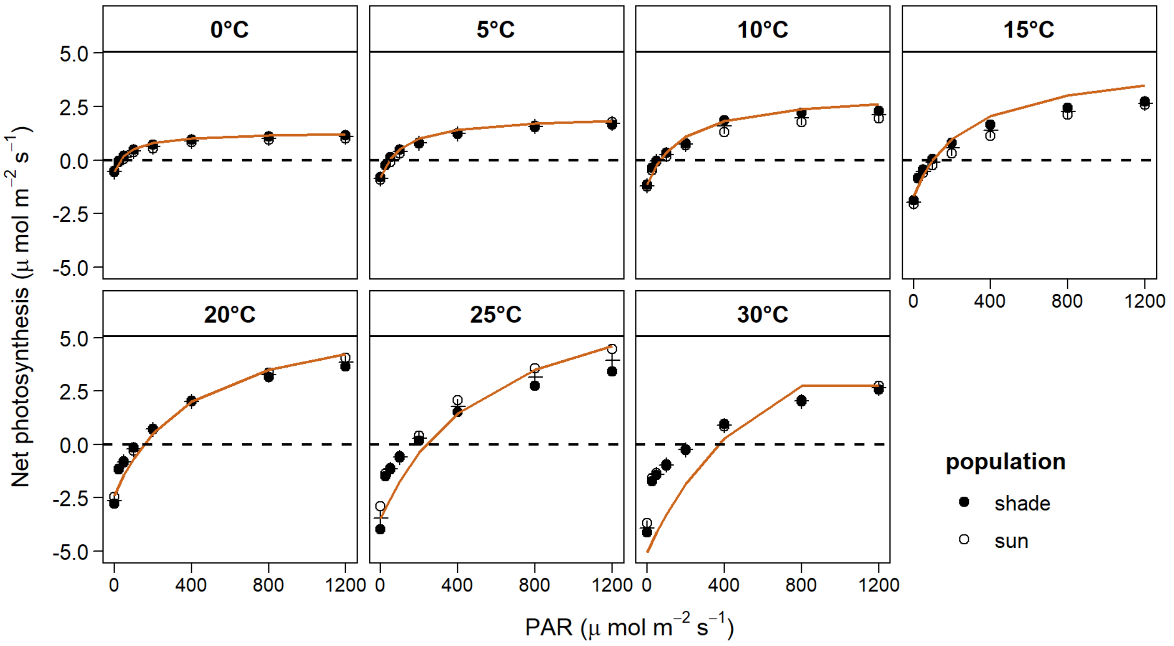


**Fig. S2** Calibration results for photosynthesis: Modeled and measured light curves of net photosynthesis at different temperatures. Points represent measured data of a sun and shade populations of *Diploschistes diacapsis* (data from Pintado et al. (2005), the cross represents the mean of sun and shade population and lines modeled values.

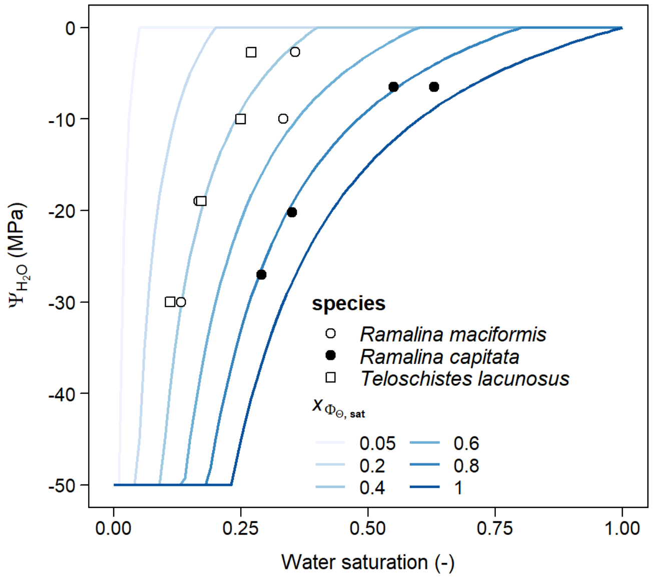


**Fig. S3** Water potential $\Psi_{H2O}$ as function of changing thallus water saturation Θ. Points represent measured relationships of the three different dryland lichen species, lines show the relationship for different values of the threshold saturation $x_{\Phi\Theta,sat}$. Ramalina capitata data from central Spain (Pintado & Sancho, 2002), and Ramalina capitata and Teloschistes lacunosus from the Negev desert (Scheidegger, Schroeter, & Frey, 1995).

**Table S2** Literature and calibrated values for parameters of *Diploschistes diacapsis* used in the simulation experiments.

| Parameter | Value | Unit | Reference | Details |
|---|---|---|---|---|
| **albedo** | 0.298 | - | (Chamizo et al., 2012) | Average reflectance of short-wave radiation (square root of sum of squares of the reflectance of light-colored lichen dominated biocrusts at every wavelength between 300 and 1100 nm) |
| **thallus height** | 1.4 | mm | (Pintado et al., 2005) | Mean thallus height (sun and shade population) of *D. diacapsis* |
| **thallus porosity** | 0.3-0.4 | - | - | Chosen such that resulting specific thallus area is between 3.6 and 4.2 $m^2$ thallus $kg^{-1}$ C |
| **water storage capacity** | 0.58 | % of dry weight | (Pintado et al., 2005) | Mean water storage capacity (sun and shade population) of *D. diacapsis* |
| **saturation at maximum activity** | 0.3 | - | (Pintado et al., 2005) | Deduced from the relation between net photosynthesis and thallus water saturation (Fig. 1b) |
| **saturation at which water potential becomes negative** | 0.05 – 1 | - | - | Unknown, range of possible values was used (Fig. S3) |
| **optimum temperature for photosynthesis** | 25 | °C | (Pintado et al., 2005) | Maximum of temperature dependent gross photosynthesis; mean of sun and shade population |

| | | | | |
|---|---|---|---|---|
| **$Q_{10}$ value of respiration** | 2.1 | - | (Pintado et al., 2005) | Mean of all $Q_{10}$ values calculated from the relationship between temperature and dark respiration with $Q10 = \frac{R1^{\frac{10}{T1-T2}}}{R2}$ with R1 and R2 being the dark respiration rates at temperature T1 and T2 respectively |
| **reference maintenance respiration** | 3.5 | μmol $m^{-2}s^{-1}$ | (Pintado et al., 2005) | Calculated from dark respiration at 10°C ($R_{spec}$), temperature ($T_{surf}$, here 10°C), optimum temperature for photosynthesis ($T_{opt}$) and the Q10 value: $$R_{main} = \frac{R_{spec}}{Q10^{\frac{T_{surf}-T_{opt}}{10}}}$$ |
| **$V_{C,max}$** | 2.1 | $s^{-1}$ | | |
| **$V_{O,max}$** | 0.3 | $s^{-1}$ | | |
| **thallus $CO_2$ diffusivity** | 0.01 | mol $m^{-}2$ $s^{-1}$ | | |
| **enzyme activation energy $K_c$** | 80000 | J $mol^{-1}$ | (Pintado et al., 2005) | Calibration using light curve (Fig. S2) |
| **enzyme activation energy $K_o$** | 55000 | J $mol^{-1}$ | | |
| **activation energy $V_m$** | 10000 | J $mol^{-1}$ | | |
| **activation energy $J_m$** | 60000 | J $mol^{-1}$ | | |
| **$x_{\Phi\Theta,sat}$ threshold saturation for water potential** | 0.05-1 | - | (Pintado & Sancho, 2002, Scheidegger et al., 1995)) | Values were chosen such that the obtained curves are within a range of relationships observed for the |

| | | | | |
|---|---|---|---|---|
| **$x_{\Psi H2O}$ shape parameter for the water content dependent water potential curve** | 15 | - | | dryland lichens *Ramalina capitata* in Central Spain *and Ramalina maciformis* and *Teloschistes lacunosu*s from the Negev desert in Israel (see Fig. S3) |

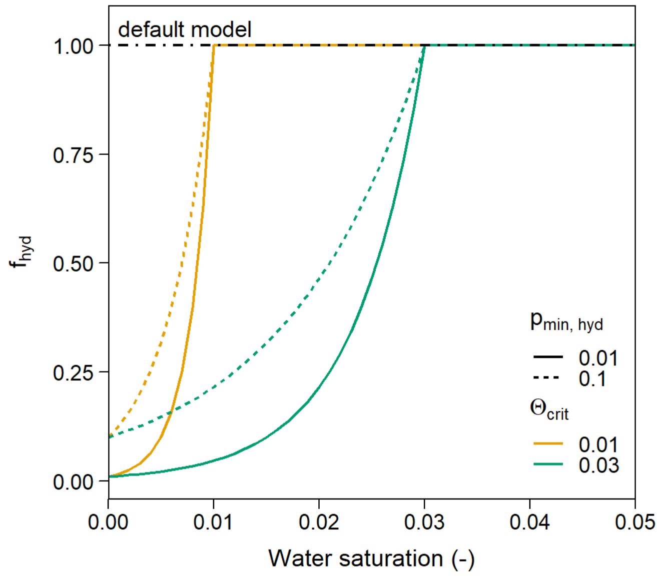


**Fig. S4** Exponential increase of $f_{hyd}$ for different parameterizations of $p_{min,hyd}$ and $\Theta_{crit}$. The default model without hydrophobicity corresponds to a value of 1 for $f_{hyd}$ independent of the current thallus water saturation $\Theta$.

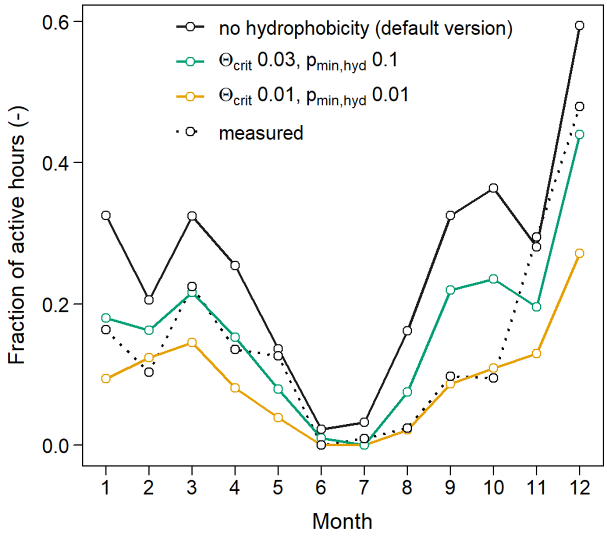


**Fig. S5** Calibration of the hydrophobicity parameters against mean monthly active time *of Diploschistes diacapsis*. For different parameterizations of the hydrophobicity function, the simulated proportion of active time (continuous line) is compared with measured activity.

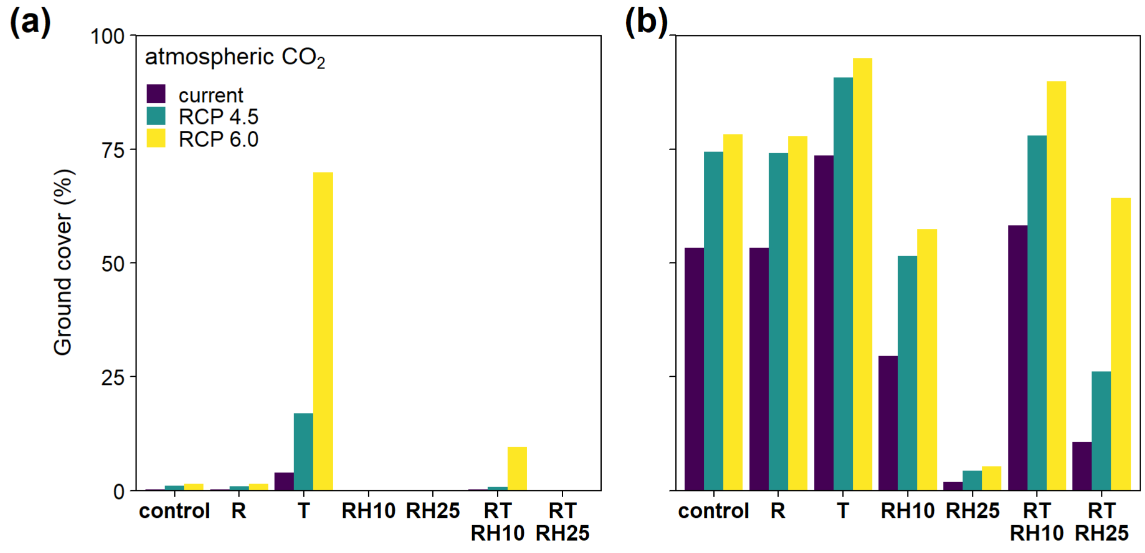


**Fig. S6** Lichen cover for different climate scenarios and hydrophobicity parameterizations. **(a)**: Hydrophobicity parameterized with $\Theta_{crit}$ = 0.01 and $p_{min,hyd}$ = 0.01, **(b)**: Default model without hydrophobicity. Note that for (a) the simulation was only run for 200 instead of 900 years because for this parameterization of hydrophobicity, the species has a 100% mortality rate after 200 years. Bars represent cover values for different atmospheric CO2 concentrations (current = 395 ppm, RCP 4.5 = 650 ppm, RCP 6.0 = 850 ppm) and climate scenarios (control = no changes in rainfall, temperature and relative humidity, R = rainfall reduction by 30%, T = temperature increase by 3 °C for current and RCP 4.5 scenario and 5 °C for RCP 6.0, RH10 = reduction of relative humidity by 10%, RH25 = reduction of relative humidity by 25%, RTRH10 = combination of R, T, and RH10 scenario, RTRH25 = combination of R, T, and RH25 scenario).

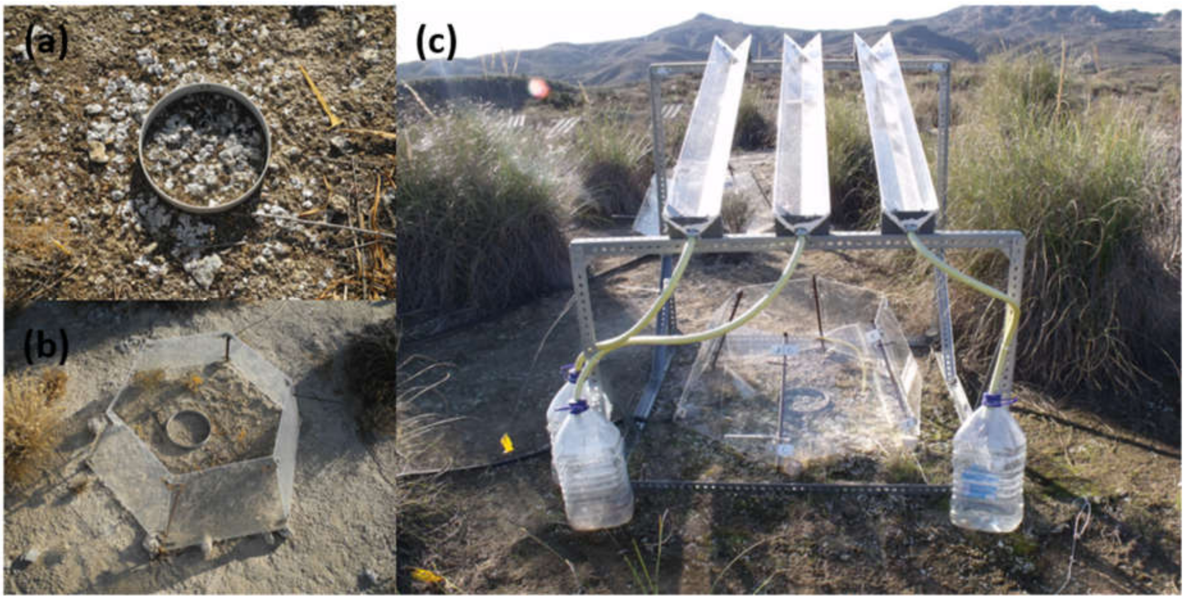


**Fig. S7** Experimental setup of the manipulative climate change experiment in Aranjuez. **(a)**: control treatment, **(b)**: warming treatment with the open top chamber, **(c)**: combination scenario with a combination of warming and rainfall exclusion with a rainfall shelter. Note: the treatment where only rainfall is reduced by a rainfall shelter is not shown here (Photos (a) and (b) by Selina Baldauf, Photo (c) by Fernando T. Maestre).

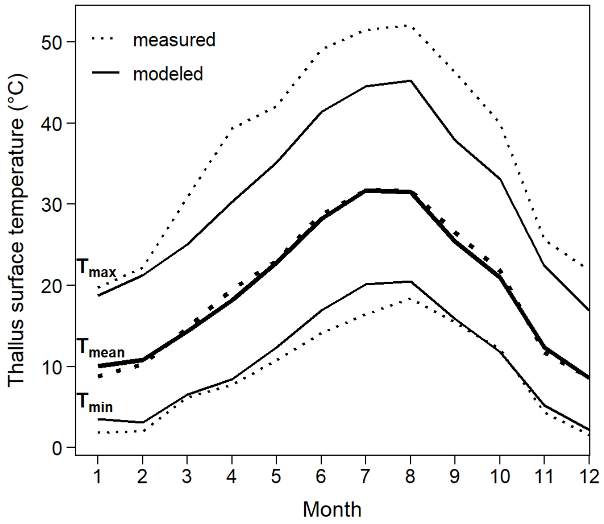


**Fig. S8** Measured and modeled mean, minimum and maximum monthly *Diploschistes diacapsis* thallus surface temperature in El Cautivo.

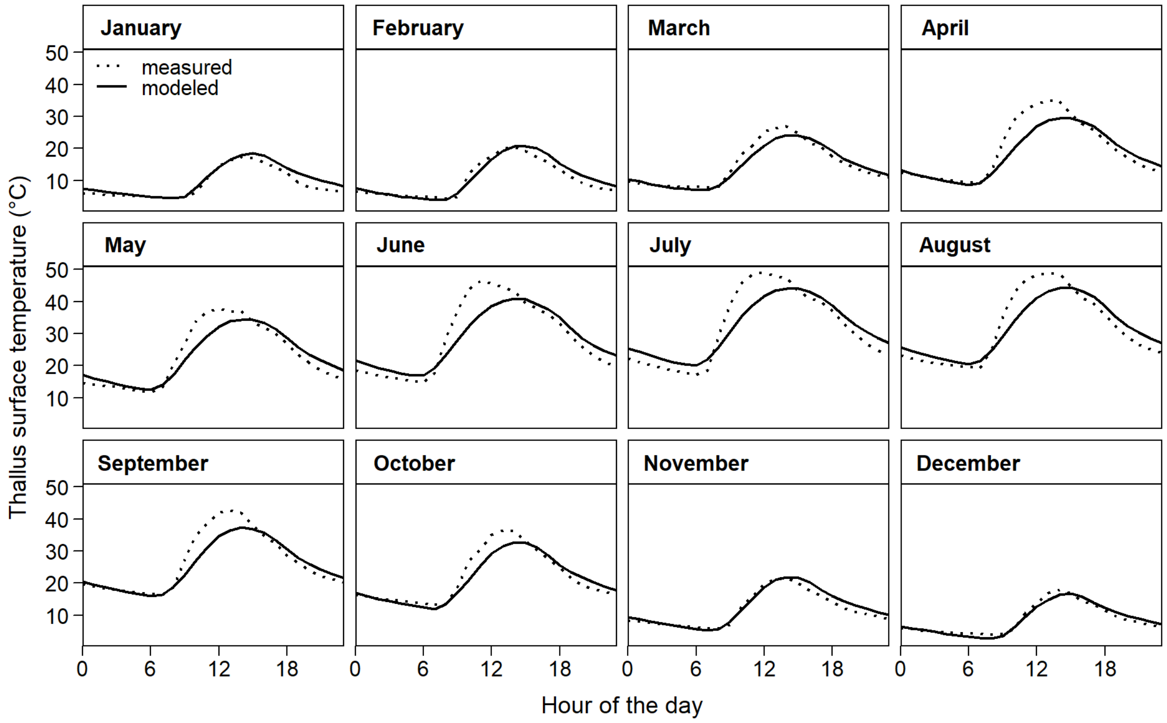


**Fig. S9** Measured and modeled hourly means of *Diploschistes diacapsis* thallus surface temperature in El Cautivo.

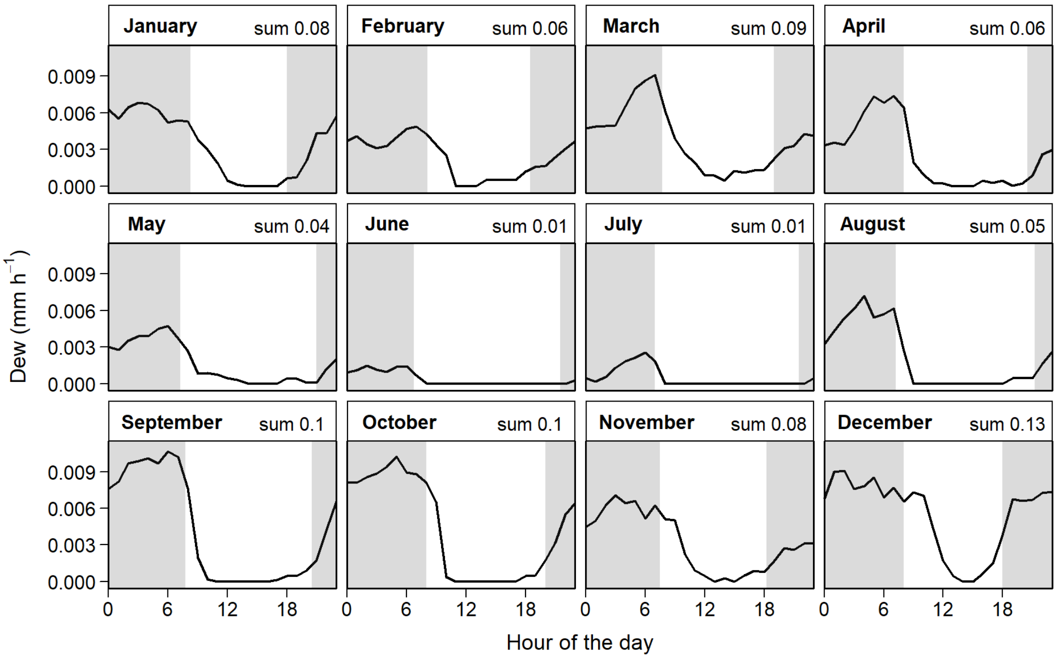


**Fig. S10** Modeled mean hourly dew rates and mean sums per day for the El Cautivo site. The grey areas in the plots indicate times before sunrise and after sunset.

**Table S2** Comparison of nights with dew input (%), total lichen active time fraction (%) and steady state cover (%) for the different climate change scenarios.

| Atmospheric $CO_2$ | Scenario[1] | Dew nights (%) | Active time (%) | Steady state cover (%) |
|---|---|---|---|---|
| current | control | 78 | 16 | 36 |
| RCP 4.5 | control | 78 | 16 | 66 |
| RCP 6.0 | control | 78 | 16 | 68 |
| current | R | 78 | 16 | 37 |
| RCP 4.5 | R | 78 | 16 | 67 |
| RCP 6.0 | R | 78 | 16 | 69 |
| current | T | 81 | 37 | 62 |
| RCP 4.5 | T | 79 | 18 | 79 |
| RCP 6.0 | T | 80 | 19 | 87 |
| current | RH10 | 67 | 11 | 8 |
| RCP 4.5 | RH10 | 67 | 11 | 23 |
| RCP 6.0 | RH10 | 67 | 11 | 28 |
| current | RH25 | 47 | 4 | 0 |
| RCP 4.5 | RH25 | 47 | 4 | 1 |
| RCP 6.0 | RH25 | 47 | 4 | 2 |
| current | RTRH10 | 73 | 25 | 31 |
| RCP 4.5 | RTRH10 | 70 | 12 | 54 |
| RCP 6.0 | RTRH10 | 72 | 13 | 72 |
| current | RTRH25 | 55 | 11 | 1 |
| RCP 4.5 | RTRH25 | 50 | 5 | 3 |
| RCP 6.0 | RTRH25 | 54 | 6 | 8 |

[1] R = rainfall reduction by 30%, T = temperature increase by 3 °C for current and RCP 4.5 scenario and 5 °C for RCP 6.0, RH10 = reduction of relative humidity by 10%, RH25 = reduction of relative humidity by 25%, RTRH10 = combination of R, T, and RH10 scenario, RTRH25 = combination of R, T, and RH25 scenario.

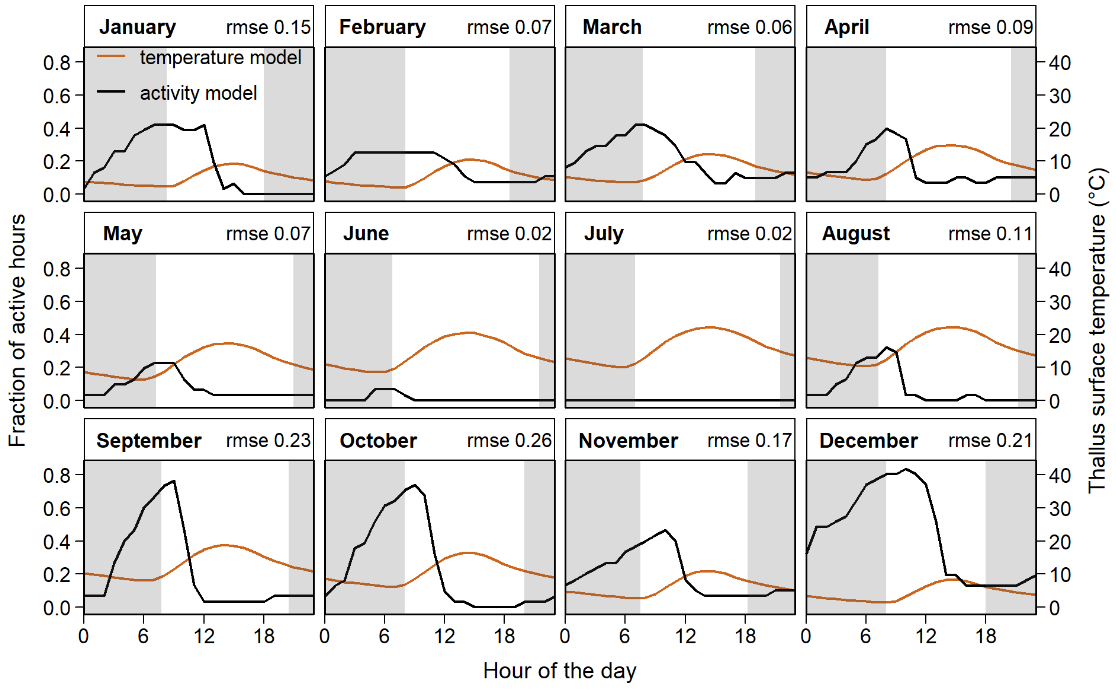


**Fig. S11** Modeled diurnal fraction of active hours in each month and modeled thallus surface temperature. The grey areas in the plots indicate times before sunrise and after sunset.
